# Non-canonical TERT function mitigates adipose tissue inflammation in obesity by modulating stem cell-macrophage communication and macrophage states

**DOI:** 10.64898/2025.12.18.694395

**Authors:** Laura Braud, Andrea Ninni, Antonio M.A. Miranda, Emilie Baudelet, Julien Vernerey, Marina Villaverde, Francesca Sciarretta, Manuel Bernabe, Léa Girondier, Ilaria Elia, Jesus Gil, Stéphane Audebert, Jean-Marc Brondello, Christophe Lachaud, Will Scott, Daniele Lettieri-Barbato, Vincent Géli

## Abstract

**Background and aims:** Obesity drives adipose tissue (AT) expansion and chronic inflammation, leading to metabolic dysfunction through adipocyte hypertrophy, impaired ASPC differentiation, immune infiltration, and fibrosis. Stromal vascular remodeling prominently features expansion of Gdf15- and Trem2-expressing lipid-associated macrophages (LAMs), which respond to adipocyte stress and act as lipid scavengers to buffer excess lipids released by adipocytes. Here, we examined macrophage reprogramming in p21^+/Tert^ and p21^+/TertCi^ mice, which express active TERT or its catalytically inactive TERT^Ci^ mutant from the endogenous *Cdkn1a* promoter.

**Results:** Following HFD exposure, conditional expression of TERT or TERT^Ci^ resulted in a pronounced downregulation of p21 in macrophage subsets, accompanied by a reduction in AT inflammation. Notably, TERT and TERT^Ci^ expression reshaped the adipose-tissue macrophage (ATM) landscape depleting Trem2+ and Gdf15+ LAMs while preserving resident macrophages. This shift was accompanied by marked suppression of the Trem2 transcriptional program and down-regulation of PPAR-γ and NR1H3 in LAMs and by impaired ASPC-LAM signaling pathways that normally drive LAM recruitment and activation. Proteomic profiling further showed that p21^+/Tert^ ASPCs secreted markedly higher levels of proteins associated with non-conventional secretion, while extracellular matrix–associated factors and cytokines/chemokines including key mediators of ASPC–LAM communication such as Ccl2, C3, and Csf1 were substantially reduced. However, only p21^+/Tert^ mice, and not p21^+/TertCi^ mice, exhibited significant metabolic improvements, indicating that macrophage remodeling alone is insufficient to restore systemic metabolic function. Consistent with this, enhanced ASPC expansion and differentiation, supporting improved adipose-tissue remodeling, was observed exclusively in p21^+/Tert^ obese mice.

**Conclusions:** TERT remodels adipose tissue immunity independently of its enzymatic activity, and TERT-driven reprogramming of the ASPC secretome may emerge as a promising strategy to combat obesity-related metabolic dysfunction.

**Highlights:**

- Conditional expression of TERT or catalytically inactive TERT^Ci^ results in the depletion Trem2+ and Gdf15+ LAMs while preserving resident macrophages
- TERT and TERT^Ci^ expression impairs ASPC-adipocyte/LAM communication pathways that normally drive LAM recruitment and activation.
- In vitro, TERT conditional expression in ASPCs promotes non-conventional protein secretion and reduces the secretion of key mediators of ASPC–LAM communication
- Only p21^+/Tert^ mice, and not p21^+/TertCi^ mice, exhibited enhanced ASPC expansion, improved adipose-tissue remodeling, and systemic metabolic benefits, demonstrating that macrophage remodeling alone is insufficient to restore metabolic function.

## Introduction

Obesity is a chronic metabolic disorder characterized by excessive adipose tissue expansion and systemic low-grade inflammation, which are closely associated with the development of insulin resistance, type 2 diabetes, non-alcoholic fatty liver disease (NAFLD), and cardiovascular disease [1]. During obesity, numerous events participate in adipose tissue dysfunction, including adipocyte hypertrophy, impaired adipose stem and progenitor cell (ASPC) differentiation capacity, pro-inflammatory immune cell infiltration and fibrosis [2]. The stromal vascular fraction (SVF) of the adipose tissue that includes preadipocytes, fibroblasts, endothelial cells, pericytes, and immune cells undergoes significant remodelling driven by changes in cellular composition, vascularization, and extracellular matrix (ECM) composition [3–6]. In particular, the proportion of macrophages in the adipose tissue (ATM) can rise from 10% to 40% in obese people [7]. Distinct ATM subsets, including lipid-associated macrophages (LAMs), have been identified, with specialized roles in lipid uptake, tissue remodeling, and inflammatory signaling [4,8]. ATMs are recruited by different factors secreted by the AT [9]. The recruited macrophages perform several functions both in health and disease, including elimination of dead adipocytes by phagocytosis, tissue homeostasis, insulin resistance, inflammation, and adipose tissue fibrosis [10]. Studies with High Fat Diet (HFD)-fed mice have demonstrated that ATMs not only increase in number but also exhibit metabolic reprogramming, including enhanced glycolysis, altered lipid handling, and senescence [4,11,12]. A specific subset of LAMs is characterized by the expression of the lipid receptor Trem2. These Trem2-positive macrophages that accumulate around lipid-laden dying adipocytes are responsible for lipid absorption [6].

Mouse models of diet-induced obesity have been instrumental in uncovering mechanisms that drive metabolic dysfunctions. In particular, studies in mice have shown that targeted inactivation of the catalytic subunit of telomerase (TERT) in adipose stem and progenitor cells (ASPCs) leads to cellular senescence, which is associated with adipocyte hypertrophy, increased inflammation, fibrosis, and systemic insulin resistance [13]. Notably, one mouse study identified a rare population of Tert-expressing adipose stem cells capable of differentiating into mature adipocytes in response to a high-fat diet (HFD) [14]. Despite these insights, TERT expression or activity in the context of obesity has been largely unexplored in both humans and mice. Importantly, numerous studies have shown that TERT exerts functions beyond its canonical role in telomere elongation [15]. Depending on the cellular context, TERT can positively modulate multiple signalling pathways, including WNT, NF-κB, and MYC [16–18]. Most recently, work using dextran sulfate sodium treated knock-in mice expressing a catalytically inactive form of Tert revealed that TERT regulates inflammatory gene expression independently of its enzymatic activity, underscoring a non-canonical pro-inflammatory role for TERT under these conditions [19].

We generated two mouse models (called p21^+/Tert^ and p21^+/TertCi^) that expresses the telomerase reverse transcriptase (TERT) or its catalytically inactive form (TERT^Ci^) under the control of the *p21*-promoter [20]. We previously demonstrated that TERT expression in the lungs of p21^+/Tert^ mice reduces p21 levels, diminishes oxidative damage, limits endothelial cell senescence, and prevents the age-associated loss of capillary density [20]. More recently, using the same p21^+/Tert^ model, we showed that conditional TERT expression ameliorates obesity-associated metabolic dysfunction, primarily by suppressing senescence-related signaling across multiple cell types and by promoting ASPC expansion and adipogenic differentiation [21]. These findings prompted us to compare the phenotypes of p21^+/Tert^ and p21^+/TertCi^ mice under conditions of HFD-induced obesity.

Our results show that both models exhibit reduced p21 expression particularly within macrophages accompanied by decreased inflammation in the AT of obese mice. Notably, our data reveal a catalytic-activity–independent role for TERT in shaping macrophage composition within AT, characterized by a reduction in Trem2-positive and Gdf15-positive LAMs and a corresponding increase in resident macrophages. We found that this shift in macrophage composition was linked to a TERT-driven alteration in the crosstalk between ASPCs and macrophages. Nevertheless, the reduction in inflammation observed in p21^+/TertCi^ mice appears insufficient to fully restore metabolic function, when compared to the improvements seen in p21^+/Tert^ mice. Our study reveals that TERT expression under the control of the p21-promoter reprograms the secretome of ASPC derived from mouse adipose tissue uncovering new therapeutic avenues.

## Results

### Both TERT and TERT^Ci^ reduce p21 expression in AT macrophages

Male p21^+/mCherry^ mice were maintained on a standard diet (SD) as controls, while p21^+/mCherry^, p21^+/Tert^, and p21^+/TertCi^ mice were fed a high-fat diet (HFD) for 8 weeks as previously described [21] (Figure 1A). As expected, HFD feeding led to a significant increase in body weight compared to SD-fed controls (Figure 1B). However, no significant differences in body weight gain were observed among the HFD-fed p21^+/-^, p21^+/Tert^, and p21^+/TertCi^ groups (Figure 1B). This weight gain was associated with increased visceral adipose tissue (AT) mass in all HFD-fed mice, with no significant variation between genotypes (Figure 1C). We previously showed that HFD-fed mice exhibited elevated p21 expression in adipose tissue macrophages which was reduced in p21^+/Tert^ mice [21]. To further characterize p21-expressing ATMs, we analyzed stromal vascular fraction (SVF) cell subsets isolated from p21^+/−^, p21^+/Tert^, and p21^+/TertCi^ mice by flow cytometry. Using established surface markers together with the validated gating strategy previously described [21], we delineated discrete SVF populations and identified the corresponding p21^+/mCherry^-expressing subsets within each (Figure 1D). Of note, HFD feeding significantly increased mCherry signal across all macrophage populations, with the strongest induction observed in macrophages expressing CD206 (mannose receptor) and CD301 (C-type lectin) (Figure 1D). CD206 and CD301 are hallmarks of alternatively activated macrophages (M2-like), which are typically associated with anti-inflammatory and tissue-repair functions such as extracellular matrix remodelling, angiogenesis, and maintenance of tissue homeostasis [22,23]. Importantly, expression of both Tert and catalytically inactive Tert^Ci^ markedly reduced HFD-induced p21 upregulation in macrophages (Figure 1E-H). This reduction in p21 expression was further confirmed in the SVF by qPCR analysis (Figure 1I), consistent with the flow cytometry findings. As previously reported [21], we observed that HFD feeding increases the fraction of short telomeres (<1 kB) in the SVF of AT (Figure 1J-K). While Tert expression prevents this telomere shortening, the catalytically inactive Tert^Ci^ had not such effect (Figure 1K-J) supporting a non-canonical role for Tert in regulating p21 expression. Taken together, these results demonstrate that Tert reduces the proportion of p21-positive macrophages in AT independently of its catalytic activity.

**Figure 1.**
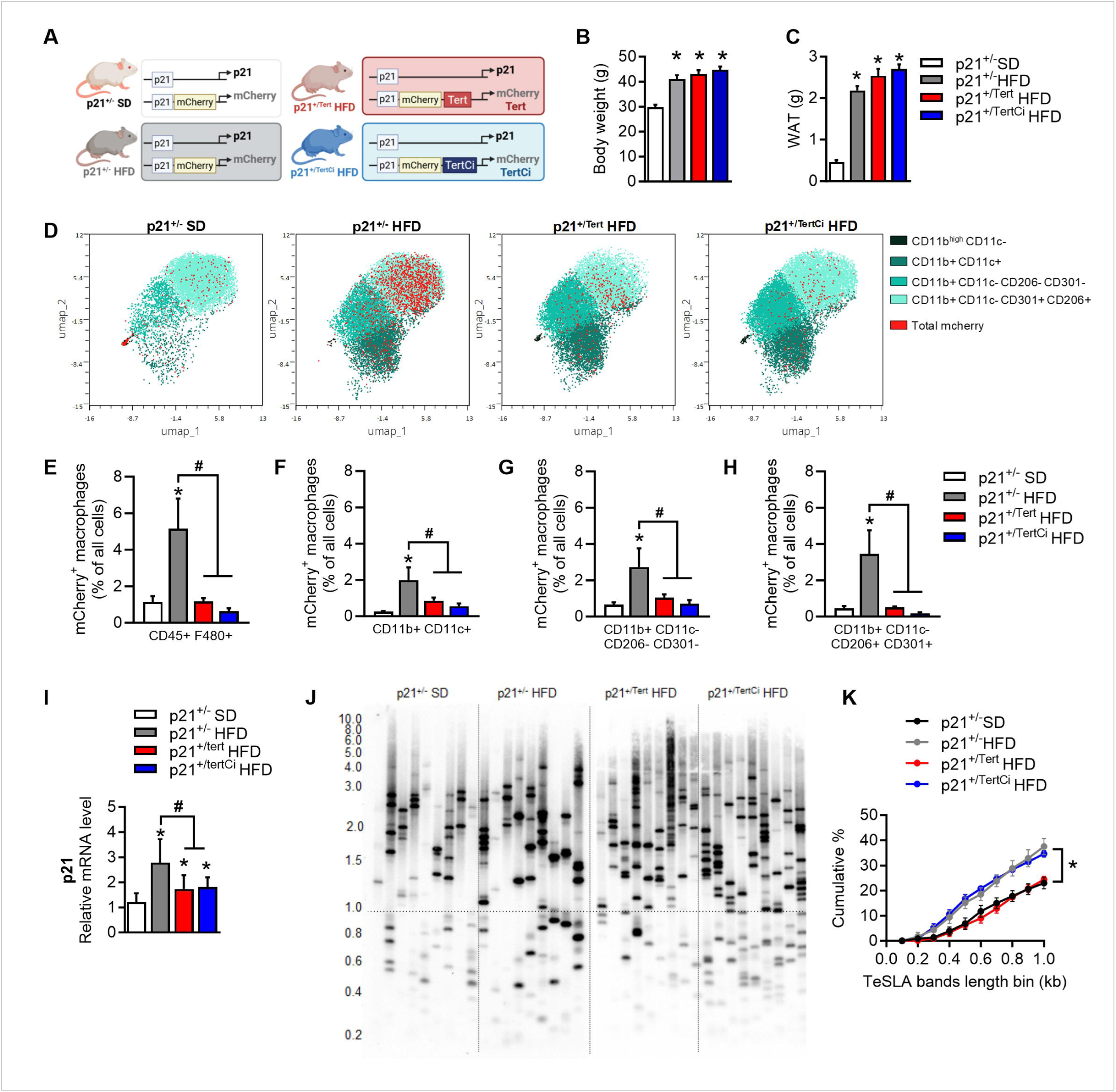
TERT and TERT^Ci^ reduce p21-expressing macrophages in the adipose tissue of obese p21^+/Tert^ HFD and p21^+/TertCi^ HFD mice. A. Schematic representation of p21^+/-^, p21^+/Tert^ and p21^+/TertCi^ mouse models B. Body weight (g) of p21^+/-^ SD, p21^+/-^ HFD, p21^+/Tert^ HFD and p21^+/TertCi^ HFD mice C. Visceral adipose tissue (AT) weight of mice at sacrifice D. UMAP clustering of macrophages and mCherry positive cells. Macrophages populations were distinguished by FACS using the markers as previously described [21] E. Percentage of mCherry positive macrophages CD45^+^ F480^+^ F. Percentage of mCherry positive macrophages CD11b^+^ CD11c^+^ G. Percentage of mCherry positive macrophages CD11b^+^ CD11c^-^ CD206^-^ CD301^-^ H. Percentage of mCherry positive macrophages CD11b^+^ CD11c^-^ CD206^+^ CD301^+^ I. p21 mRNA levels was measured by RT-qPCR in stromal vascular fraction (SVF) J. Representative image of short telomere fraction in the SVF cells isolated from the indicated mice by Telomere Shortest Length Assay (TeSLA) K. Analysis of the short telomere fraction in the SVF cells isolated from the indicated mice by Telomere Shortest Length Assay (TeSLA) [21]. *p<0.05 vs. p21^+/-^ SD mice (white bars), ^#^p<0.05 vs. p21^+/-^ HFD (grey bars). n=6 mice/group. Student’s t test or one-way ANOVA with Fisher multiple comparison test.

### Expression of either TERT or TERT^Ci^ attenuates inflammation

To investigate the broader transcriptional impact of conditional Tert and Tert^Ci^ expression as well as and the reduction of p21-positive cells, we performed RNA sequencing (RNA-seq) on SVF samples from p21^+/-^, p21^+/Tert^, and p21^+/TertCi^ mice fed an HFD, alongside p21^+/-^ SD mice (Figure 2) (Dataset 1).

**Figure 2.**
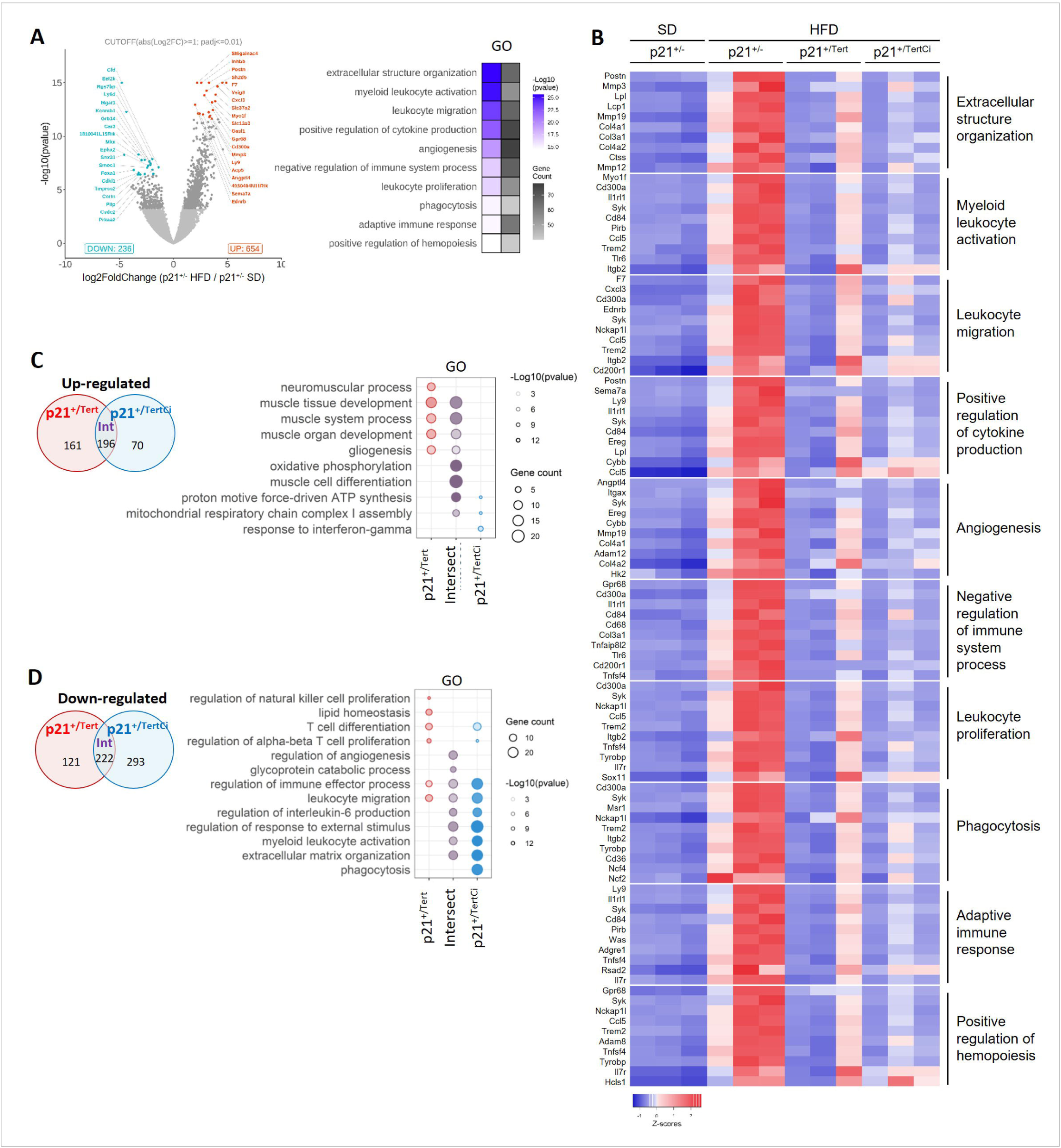
Transcriptomic profiles of the adipose tissue SVF from obese p21^+/Tert^ HFD and p21^+/TertCi^ HFD mice. RNA was purified from SVF isolated from three independent mice of the indicated genotypes. A. Volcano plots indicating the number of significantly (<0.05) differentially expressed between p21^+/-^SD and p21^+/-^ HFD mice and associated Gene Ontology (GO) analysis revealing enrichment of genes involved in the indicated pathways. B. Heatmap of differentially expressed genes in the SVF of the indicated mice. Genes belong to the functional categories shown on in Figure 2A. C. Venn diagram indicating the number of significantly (<0.05) up-regulated between p21^+/-^ and p21^+/Tert^ or p21^+/TertCi^ HFD mice. GO enrichment analysis of up-regulated genes in p21^+/Tert^ HFD only, p21^+/TertCi^ HFD only and both, compared to p21^+/-^ HFD mice. D. Venn diagram indicating the number of significantly (<0.05) down-regulated between p21+/- and p21+/Tert or p21+/TertCi HFD mice. Gene ontology (GO) enrichment analysis of down-regulated genes in p21^+/Tert^ HFD only, p21^+/TertCi^ HFD only and both, compared to p21^+/-^ HFD mice. n=3 mice/group

We first compared gene expression changes between standard diet (SD)- and (HFD)-fed p21^+/⁻^ mice and identified 654 genes upregulated and 236 downregulated in response to HFD (Figure 2A, left panel), highlighting the high plasticity of AT [24]. The volcano plot shows the top 20 most significantly up- and down-regulated genes. Gene Ontology (GO) analysis revealed 54 enriched biological processes (adjusted p-values ranging from 10⁻²² to 0.0025) with the top 10 pathways along including extracellular matrix organization, cytokine secretion, immune remodelling, and angiogenesis, key processes linked to obesity (Figure 2A, right panel). We focused on the 10 most significantly enriched biological processes in p21^+/-^ mice fed a HFD compared to those on SD. We then assessed how these processes were regulated in p21^+/Tert^, and p21^+/TertCi^ fed a HFD relative to p21^+/-^ mice HFD-fed controls. As shown Figure 2B, the heatmap reveals that the upregulation of genes involved in extracellular structure organization, immune-related processes, and angiogenesis, was attenuated in the SVF of both p21^+/Tert^ and p21^+/TertCi^ mice. Expression of Tert and Tert^Ci^ driven by the p21-promoter markedly suppresses the overexpression of genes associated with inflammation (Figure 2B). To further dissect the shared and distinct effect of Tert, we performed a Venn-diagram analysis (Figure 2C-D). We found that Tert and Tert^Ci^ expression under HFD conditions led to the share upregulation of 196 genes and downregulation of 222 genes. This overlap suggests that the majority of transcriptional changes are likely mediated by a non-canonical function of Tert (Figure 2C-D). Notably, we observed a shared down-regulation of genes associated with cytokine and inflammatory pathways in the SVF of HFD-fed p21^+/Tert^ and p21^+/TertCi^ mice. Conversely, both genotypes exhibited a common up-regulation of genes linked to mitochondrial oxidative phosphorylation (Figure 2C-D). However, each genotype showed distinct gene expression profiles, with subsets of genes uniquely regulated in p21^+/Tert^ or p21^+/TertCi^ HFD fed mice (Figure 2C-D). Volcano plots (Suppl Fig 1A-B) highlight the top 20 up- and downregulated genes in each comparison along with their associated GO terms. Of note, we observed an upregulation of genes associated with hormone regulated processes and muscle cell differentiation in the SVF of p21^+/Tert^ mice (Suppl Fig 1A-B).

### TERT’s non-canonical function reshapes macrophage landscape in the AT of obese mice

Given that bulk RNA-seq analysis of the SVF revealed attenuated HFD-induced inflammation (Figure 2), we next investigated whether Tert and Tert^Ci^ modulate immune cell dynamics in AT of HFD-fed mice. To this end, we analysed single-nucleus RNA-seq (snRNA-seq) datasets from adipose tissue of both p21^+/Tert^ and p21^+/TertCi^ mice. The snRNAseq analysis was conducted following the same methodology as described in Braud et al (2025) [21], resulting in the identification of 39,167 high-quality cells (Dataset 2). Subsequent clustering and annotation identified ten distinct cell clusters, consistent with findings with our previous analyses (Figure 3A). The largest immune cell population identified was the Mononuclear Phagocyte (MPC) cluster (n=20,625), making it the most prominent cluster across all samples (Figure 3A). This population was particularly expanded in the p21^+/-^ mice fed a HFD (Figure 3A). However, the increase in MPCs was attenuated in both the p21^+/Tert^ and p21^+/TertCi^ mice compared to HFD fed p21^+/-^mice (Figure 3A).

**Figure 3.**
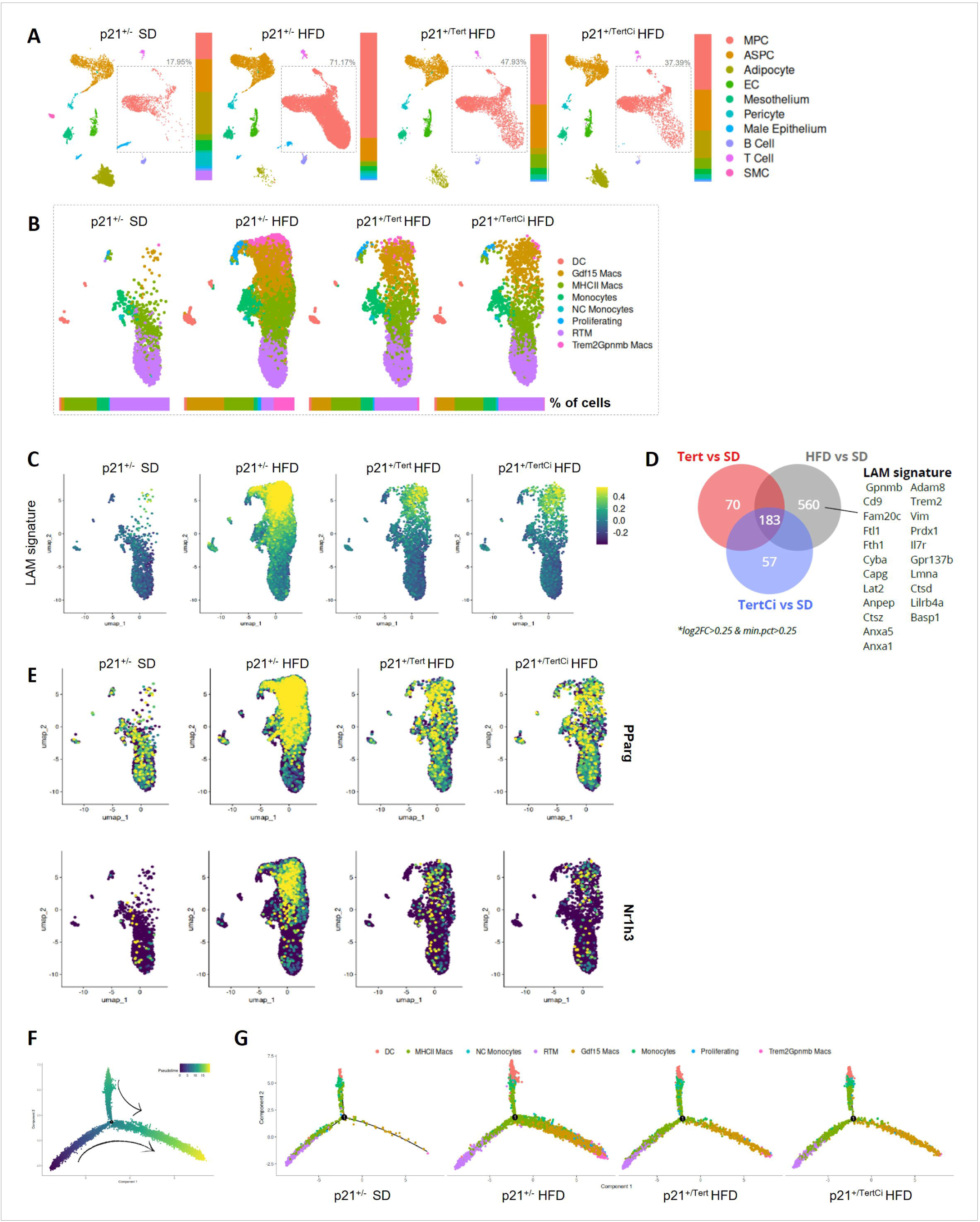
Tert’s non-canonical function reshapes macrophage landscape of obese AT. snRNA-seq was conducted on nuclei isolated from snap-frozen visceral adipose tissue collected from two independent mice per indicated genotype, and the bioinformatic analysis method is detailed in the Materials and Methods section. A. UMAP of all cell populations with integration of p21+/- lean, p21+/-, p21+/Tert and p21+/TertCi obese mice B. UMAP of macrophages subpopulations C. LAM signature (score based on the expression of established LAM-associated genes) across macrophage subclusters D. Venn diagram of the shared and unique differentially expressed genes (DEGs) in adipose-tissue macrophages from SD p21+/⁻ mice compared with HFD p21^+/⁻^, p21^+/Tert^, and p21^+/TertCi^ mice. The indicated LAM signature is present in macrophages from HFD p21^+/⁻^ adipose-tissue macrophages. E. UMAP showing PParg and Nr1h3 expression across the identified macrophage subclusters F. Trajectory analysis across the identified macrophage subclusters G. Trajectory analysis of the MPC subclusters in the different group of mice

We focussed our subsequent analysis on the MPC cluster given its central role in AT inflammation. To enable accurate and high-resolution annotation of macrophage subclusters, we first established a comprehensive macrophage annotation reference. For this, we utilized a publicly available scRNA-seq dataset derived from WAT of mice fed with a HFD for 8 and 14 weeks [25] (Suppl Fig 2). Macrophages were extracted from this dataset, and 8 distinct myeloid subclusters were defined using established marker genes from the literature (Suppl Fig 2A-B). These clusters comprised two monocyte populations (Monocytes and Non-classical Monocytes) and five macrophage populations (MHCII macs, Trem2^+^/Gpnmb^+^ macs, Gf15+ macs, resident tissue macrophages (RTM) (Suppl Fig 2A-B). The Trem2^+^/Gpnmb^+^ and Gdf15+ macrophage clusters are well-established in literature as Lipid-Associated Macrophages (LAMs) (Suppl Fig 2A).

Supervised clustering and annotation of macrophages in the four experimental groups revealed distinct patterns across the 4 models (Figure 3B) (Dataset 3). In the p21^+/-^ mice fed with a HFD, there was a marked expansion of MPC, primarily driven by the emergence of LAM clusters, particularly the Trem2+/Gpnmb+ macrophages and a significant increase in Gdf15-expressing population (Figure 3B). In sharp contrast, both p21^+/Tert^ and p21^+/TertCi^ models exhibited a markedly attenuated expansion of these clusters (Figure 3B). While they showed a moderate increase in Gdf15 macrophages, the appearance of the Trem2^+^/Gpnmb^+^ macrophage population was almost abolished (Figure 3B). To assess whether these macrophages exhibited diminished LAM characteristics, we calculated the LAM signature, a composite score based on the expression of established LAM-associated genes, as described previously [26]. As expected, the HFD p21^+/-^ showed a strong expression of this signature across both LAM clusters, with the highest levels observed in the Trem2+/Gpnmb+ cluster (Figure 3C). In contrast, expression of the LAM signature was markedly diminished in macrophages both p21^+/Tert^ and p21^+/TertCi^ models, indicating a clear impairment in the acquisition of the LAM phenotype under HFD (Figure 3C-D). Given that PPAR-γ and NR1H3 are key regulators of the LAM transcriptional program [27], we examined the expression of these transcription factors in LAM across the different experimental contexts (Figure 3E) (Dataset 3). Both factors were down-regulated p21^+/Tert^ and p21^+/TertCi^ macrophages (Figure 3E) consistent with the decreased expression of the LAM signature observed in these models (Figure 3C). Furthermore, this result was independently supported by a separate atherospectrum analysis focused on foamy characteristics, which also demonstrated a profound reduction in the macrophage foamy feature within p21^+/Tert^ and p21^+/TertCi^ mice (Suppl Fig 3A-B).

Collectively, these findings suggest either an impaired differentiation of monocyte and polarization of resident macrophages to LAMs in the p21^+/Tert^ and p21^+/TertCi^, or reduced recruitment of monocytes to AT which would otherwise give rise to LAMs. To further resolve the cell-state dynamics on MPC, we used Monocle-2 R package (v2.32.0) [28], which reconstructs transcriptional trajectories and infers changes along pseudotime (Suppl Fig 3C-D). This analysis revealed two distinct differentiation branches: one originating from resident macrophages and the other from monocytes, that converge as they differentiate into both LAM-associated clusters (Figure 3F). Notably, the LAMs (Trem2^+^/Gpnmb^+^ cells) and Gdf15 cluster appears to represent a terminal differentiation state (Figure 3F-G). Further interrogation of the inferred trajectories specifically reveals that monocytes are unable to complete the transition into the terminal Trem2^+^/Gpnmb^+^ and Gdf15 macrophage population, underscoring a defect in this differentiation pathway (Figure 3G). Together, these data indicate that telomerase expression, whether active or catalytically inactive, impairs the normal progression of monocyte- and resident-derived macrophages toward LAMs and Gdf15 clusters, likely by blunting the induction of key transcriptional regulators such as PPAR-γ and NR1H3.

### TERT and TERT^Ci^ limit the communication between ASPC/adipocytes and LAMs

Committed ASPCs in visceral AT have been identified as the initial source of obesity-induced Ccl2, a chemokine that recruits pro-inflammatory macrophages [29], highlighting a key communication axis between ASPCs/adipocytes and macrophages in obesity. To further investigate this, we analyzed cell-cell communication probabilities between adipocytes and ASPCs, Trem2^+^ and Gdf15^+^ macrophages in HFD-fed p21^+/mCherry^, p21^+/Tert^ and p21^+/TertCi^ mice (Figure 4).

**Figure 4.**
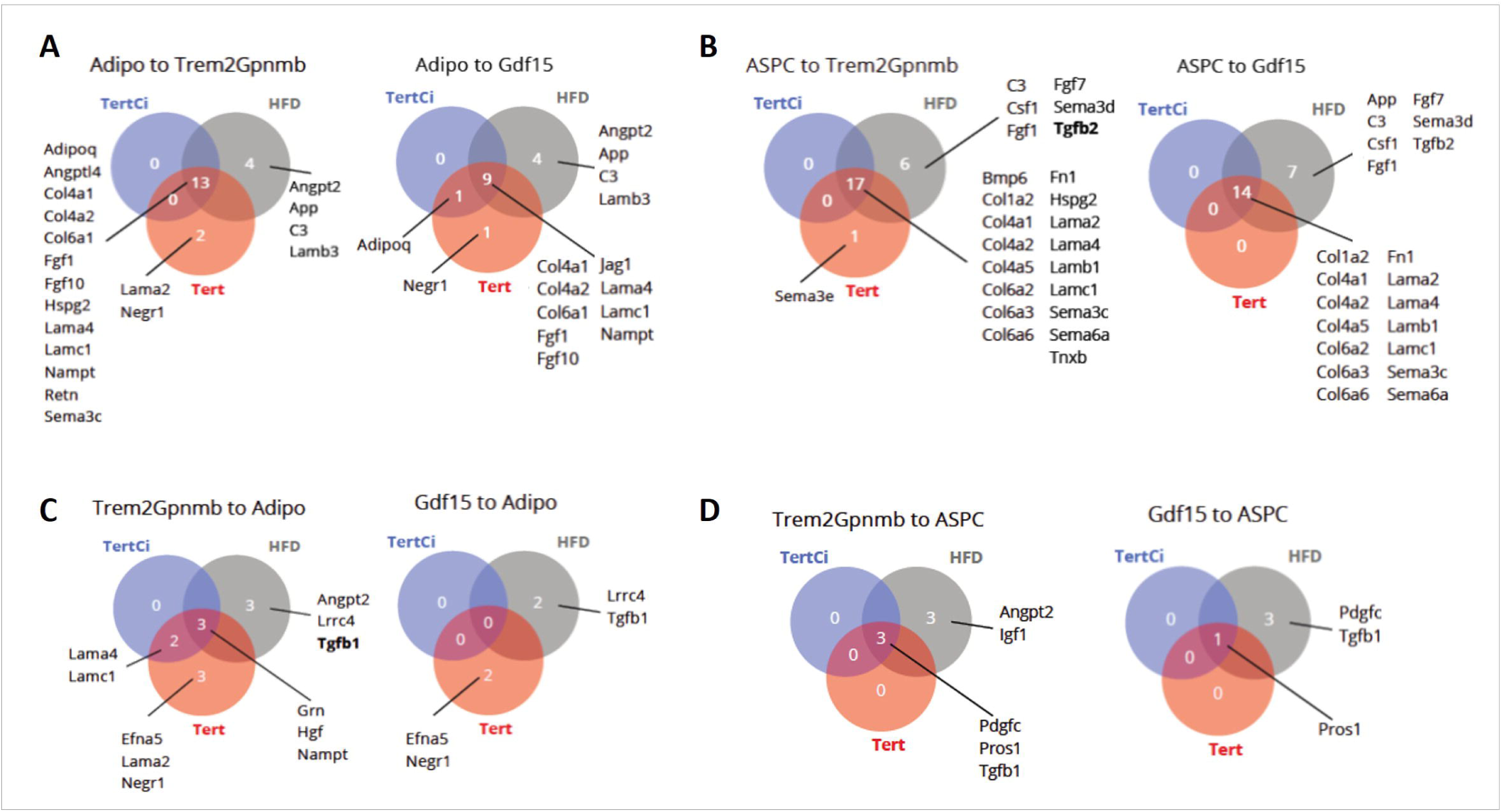
Communication probabilities between ASPC-derived adipocytes and macrophages in adipose tissue (AT) from obese p21^+/-^, p21^+/Tert^ and p21^+/TertCi^ obese mice. A. Communication probabilities from adipocytes to Trem2+ and Gdf15+ macrophages. B. Communication probabilities from ASPCs to Trem2+ and Gdf15+ macrophages. C. Communication probabilities from Trem2+ and Gdf15+ macrophages to adipocytes. D. Communication probabilities from Trem2+ and Gdf15+ macrophages to ASPCs. Communication probabilities between cell types were computed using the mass action model based on the expression of over-expressed ligand-receptor pairs.

Interestingly, ASPCs and adipocytes engaged in a high number of interactions with LAMs, as indicated by the numerous ligand–receptor pairs identified (Figure 4A-B). In contrast, Trem2+ and Gdf15+ macrophages showed weak or minimal signalling toward ASPCs and adipocytes (Figure 4C-D). Consistent with the high sender activity observed in ASPCs and adipocytes, multiple signalling interactions were conserved across the AT of all HFD-fed mouse models (Figure 4A-B). Importantly, several interactions present in the AT of p21^+/mCherry^ obese mice were absent in both p21^+/Tert^ and p21^+/TertCi^ obese mice. In particular, communication between ASPCs and Trem2+ and Gdf15+ mediated by C3, Csf1, Fgf1, Fgf7, Tgfb2, and Semad3, molecules commonly associated with inflammatory and remodelling signals, occurred only in p21^+/mCherry^ obese mice. This is consistent with the reduced inflammatory state of their adipose tissue.

### TERT expression reshapes the secretome of p21^+/Tert^ ASPCs

To validate the differences in ASPCs-LAM, we sought to compare the secretory profile of ASPCs isolated from the AT of WT and p21^+/Tert^ mice. Two pooled ASPCs preparations, each from two mice per genotype, were cultured in 10% FCS under normoxia. At ∼70% confluence, cells were left untreated or stimulated with IFN-γ (20 ng/ml) and TNF-α (10 ng/ml) for 36 hours, followed by 24 hours of serum deprivation before collecting conditioned supernatants (see Methods). Tert expression in untreated and cytokine-treated WT and p21^+/Tert^ ASPCs was assessed by RT-qPCR on harvested cells. Tert was detected exclusively in p21^+/Tert^ ASPCs and remained unchanged following IFN-γ and TNF-α stimulation (Suppl Fig 5).

Conditioned supernatants (secretomes) were analyzed for their proteomic composition using mass spectrometry. We first compared the secretory pattern of untreated WT and p21^+/Tert^ ASPCs and identified 246 proteins significantly enriched (upregulated) in the p21^+/Tert^ ASPCs supernatant compared to WT (Dataset 4). Many of these enriched proteins are involved in vesicle trafficking and endosomal transport (Figure 5A), including Rab21, Rab5a, Rab23, Sec22b, Sec23b, Sec24a, Copg2, Scfd1, Vps16, Vps25, Lamp2, Snx27, Snx18, Ap3d1, Ap3b1, and Cln5. Several members of the Cullin family (Cul1–Cul5), which serve as scaffolds for Cullin-RING E3 ubiquitin ligase complexes, were also elevated, along with the E3 ubiquitin ligase UBR4 and the autophagy cargo receptor Sequestosome-1 (SQSTM1). Notably, among the most highly enriched proteins in the p21^+/Tert^ ASPCs secretome were O-GlcNAc transferase (OGT) and Vanin1 (Vnn1) (Figure 5C). OGT catalyzes the addition of N-acetylglucosamine (GlcNAc) to serine/threonine residues and has been reported to modify numerous proteins involved in vesicle trafficking.

**Figure 5.**
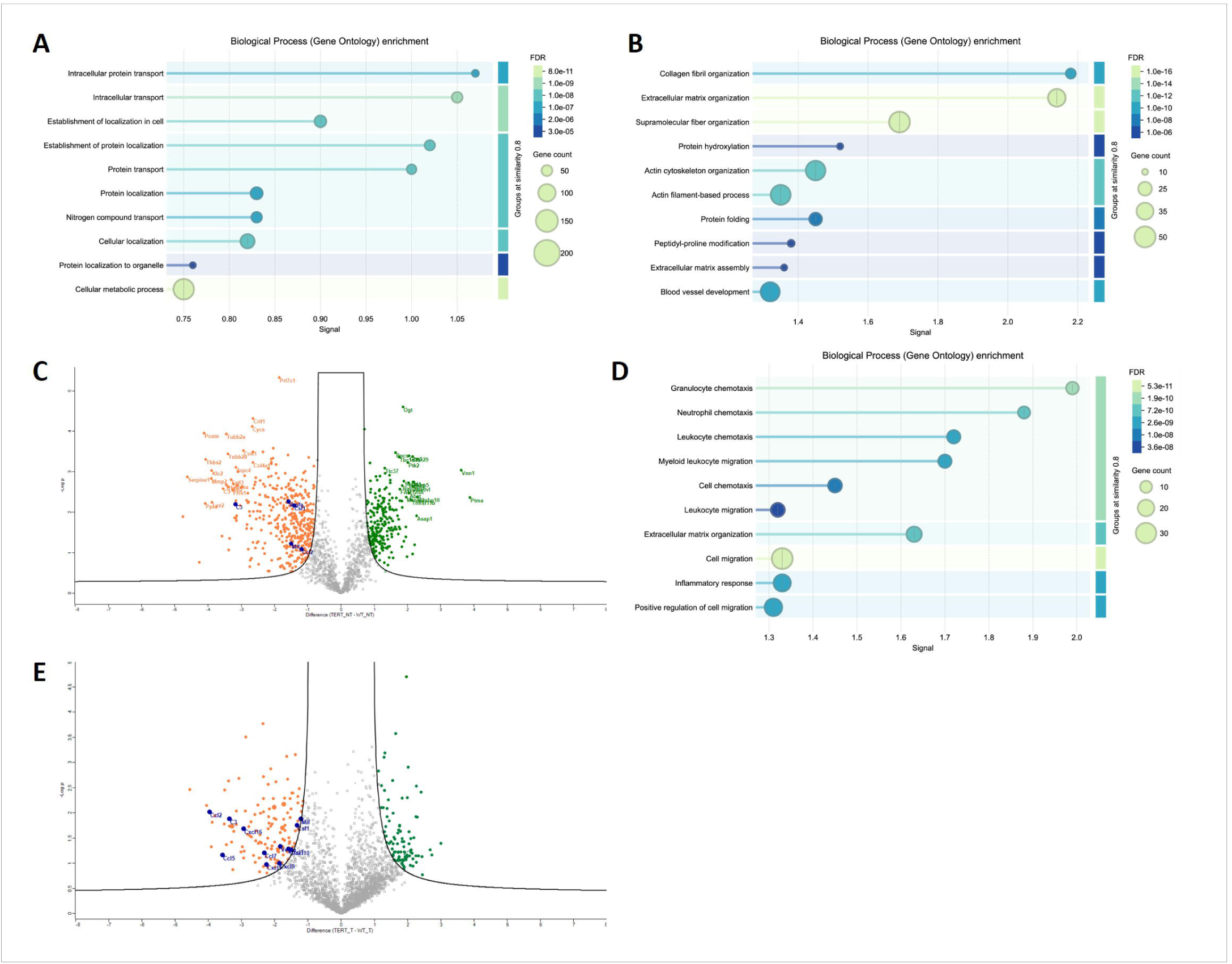
Mass spectrometry identification and analysis of secreted factors from p21+/Tert ASPCs. Supernatants from adipose stem/progenitor cells (ASPCs) of WT and p21^+/Tert^ mice were collected, and their protein content was analyzed by mass spectrometry (Dataset 4). A. Gene ontology (GO) enrichment analysis of proteins enriched in the supernatant of p21+/Tert ASPCs. B. GO enrichment analysis of proteins depleted in the supernatant of p21+/Tert ASPCs. C. Volcano plot depicting significant (<0.05) differentially represented proteins in the supernatants of WT and p21+/Tert ASPCs. Green, enriched in WT supernatants; orange, depleted in p21+/Tert supernatants. Blue dots indicate C3, CSF1, CCL2, VEGFA, and MIF. D. GO enrichment analysis of proteins depleted in the supernatant of IFN-γ and TNF-α stimulated p21+/Tert ASPCs. E. Volcano plot depicting significant (<0.05) differentially represented proteins in the supernatants of IFN-γ and TNF-α treated WT and p21+/Tert ASPCs. Blue dots indicate C3, CSF1, CCL2, MIF, FN1 and LAMC1.

We next examined the 419 proteins reduced in the p21^+/Tert^ ASPCs secretome compared to WT untreated (Dataset 4). Many are key ECM components or regulators (Figure 5B), including collagens (Col1a1, Col5a1, Col11a1), collagen-modifying enzymes (Plod1–3), ECM remodeling factors (Ecm1, Mmp2, Mmp14, Timp1, Timp2, Fn1, Lamb1), integrins, and adhesion molecules (Itga1, Itga5, Itgb5, Cdh2, Cdh11, Cd248), indicating a coordinated downregulation of ECM maintenance and cell–matrix interactions (Figure 5B). Interestingly, several underrepresented proteins (C3, Csf1, Ccl2, Vegfa, and MIF) are linked to inflammatory signaling (Figure 5C). In particular, C3 and Csf1 were found expressed in ASPCs to LAM communication analysis in the AT of WT HFD mice, but this interaction was not detected in p21^+/Tert^ mice (Figure 4B). Many proteins involved ASPC to LAM communication (Fn1, Lama2, Lamb1, Lamc1, Col4a1, Col4a2) (Figure 4B) were also reduced in the p21^+/Tert^ ASPCs secretome (Dataset 4). ApoE, a Trem2-associated protein transcriptionally repressed in ASPCs and macrophages from p21^+/Tert^ and p21^+/TertCi^ obese mice, was similarly decreased. Together, these results indicate a broad suppression of ECM and pro-inflammatory signalling in p21^+/Tert^ ASPCs.

We next compared the secretomes of IFN-γ- and TNF-α–treated WT and p21^+/Tert^ ASPCs. Among the 109 proteins that were overrepresented in the secretome p21^+/Tert^ ASPCs (Dataset 4), Annexins (Anxa2, Anxa3, Anxa4, and Anxa5), which participate in extracellular vesicle (EV) biogenesis and contribute to membrane organization, trafficking, and repair, were enriched in the secretome of p21^+/Tert^ ASPCs (Dataset 4, Figure 5E). Consistent with this observation, components of the caveolar machinery (Cav1, Cavin1, Cavin2) as well as the lysosomal membrane proteins Lamp1 and Lamp2 were also detected at higher levels in the supernatant of p21^+/Tert^ ASPCs. These findings further support the notion that TERT modulates the secretory properties of ASPCs. In addition, several proteins associated with stemness maintenance including Aldh1a1, Arhgef12, Cul4b, Lrrc40, Pik3cb, and Sod1 were more abundant in the p21^+/Tert^ ASPCs secretome (Dataset 4).

We then examined factors depleted in the secretome of p21^+/Tert^ ASPCs and identified 140 proteins significantly underrepresented relative to WT controls (Dataset 4). As observed in untreated ASPCs, proteins involved in extracellular matrix organization such as Lamb1, Lamc1, Fn1, Nid1, Mmp2, Mmp3, and Col5a1 were reduced in the supernatants of cytokine-treated p21^+/Tert^ ASPCs. GO term analysis showed that these proteins are largely associated with immune cell recruitment and migration (Figure 5D). Notably, Ccl2, Csf1, C3, and Mif were underrepresented in the p21^+/Tert^ ASPCs secretome consistent with what we previously observed in untreated cells (Dataset 4, Figure 5E). Proteomic profiling further revealed that p21^+/Tert^ ASPCs secreted markedly lower levels of cytokines/chemokines associated with leukocyte chemotaxis, myeloid cell migration, and macrophage activation, including Vegfa, Ccl2, Ccl5, Ccl7, Cxcl1, Cxcl5, Cxcl9, Cxcl10, and Cxcl16 (Dataset 4, Figure 5E). Notably, whereas cytosolic Sod1 was also enriched in the treated p21^+/Tert^ ASPCs secretome, several other oxidative stress–related proteins including Sod2, Prdx5, Prdx3, Gpx7, and Txnl1 were found at reduced levels (Dataset 4).

Together, these results indicate that IFN-γ- and TNF-α–treated p21^+/Tert^ ASPCs exhibit impaired expression or secretion of multiple immune-modulatory cytokines/chemokines.

### Bone Marrow Derived Macrophages (BMDMs) from p21^+/Tert^ and p21^+/TertCi^ obese mice exhibit activation of the Akt signaling pathway

To further characterized the potential biological pathways involved in the downregulation of inflammatory profile of the macrophages, we collected bone marrow derived macrophages (BMDMs) from SD-fed mice p21^+/-^, and obese p21^+/-^, p21^+/Tert^, and p21^+/TertCi^ mice. Because we previously observed AKT activation in ASPCs [21], we examined whether a similar effect occurred in BMDMs. We examined Akt activity and FoxO1/FoxO3 protein expression in BMDMs by western blot, factors previously reported to influence macrophage polarization [30,31]. In BMDMs from p21^+/-^ mice fed a HFD, we observed a slight (not significant) reduction in phosphorylated Akt (Akt-p) (Figure 6A-B) accompanied by a modest increase of FoxO1 and FoxO3 protein levels compared SD-fed p21^+/-^ controls (Figure 6C-D). This pattern aligns with previous reports showing that reduced Akt activity and increased FoxO1/FoxO3 expression contribute to macrophage polarization toward a pro-inflammatory phenotype [30,31]. In contrast, BMDMs derived from p21^+/Tert^ and p21^+/TertCi^ obese mice, showed increase Akt-p levels along with a significant reduction of FoxO3 and a more modest decrease of FoxO1 compared to obese p21^+/-^controls (Figure 6C-D). Gene expression analysis showed an upregulation of inflammation-associated genes (IL6, TNFa) in BMDMs isolated from obese p21^+/-^ mice, whereas their expression was significantly reduced in BMDMs isolated from both p21^+/Tert^ and p21^+/TertCi^ obese mice (Figure 6E). These results indicate that TERT, independently of its catalytic activity, helps mitigate inflammation in AT by modulating the pro-inflammatory profile of macrophages.

**Figure 6.**
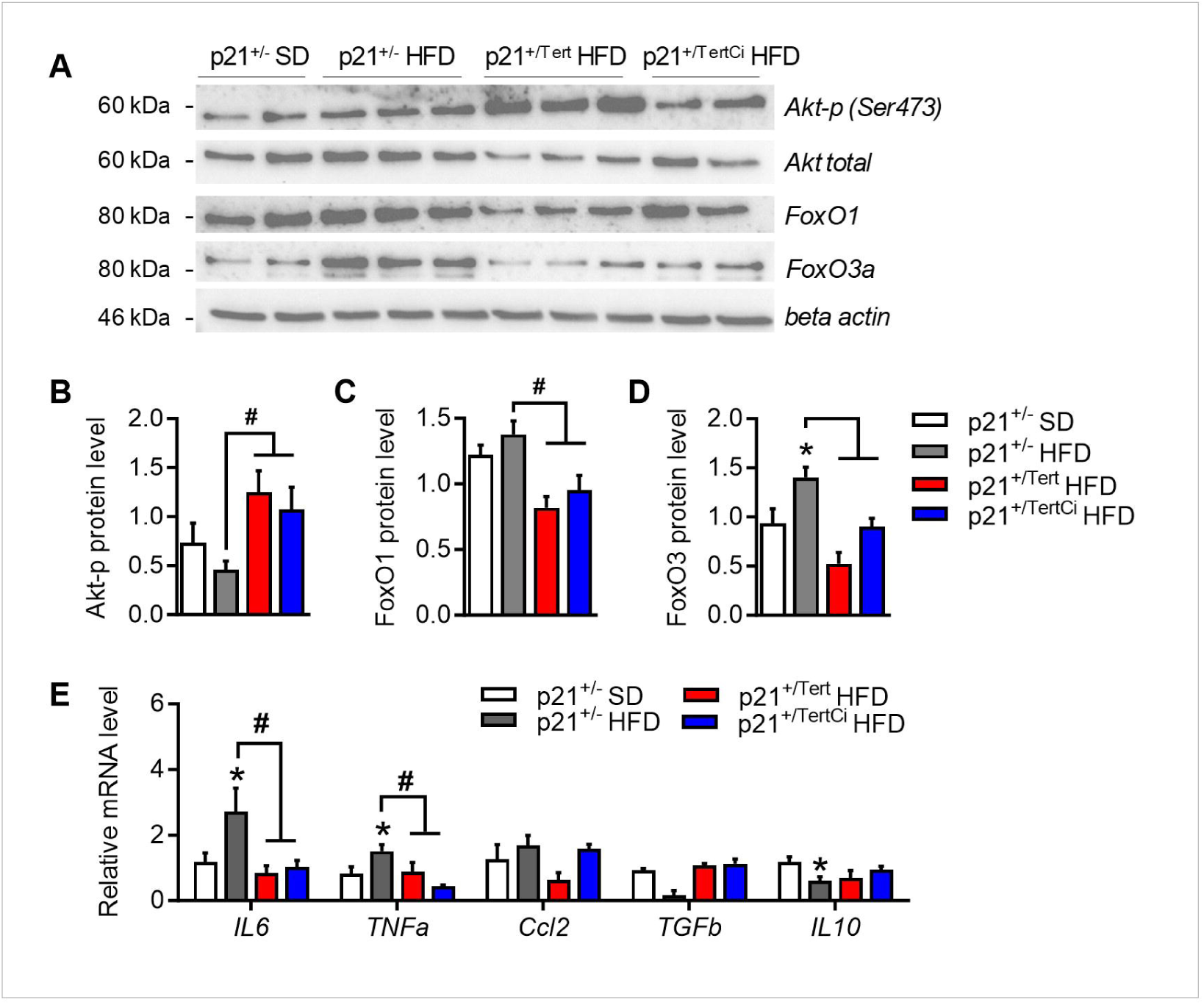
Characterization of Bone Marrow Derived Macrophages (BMDM) F. BMDM were evaluated for protein expression of total Akt, phosphorylated Akt, FoxO1 and FoxO3 assessed by Western blot G. Quantification of protein level of ratio total Akt : phosphorylated Akt(ser473) H. Quantification of protein level of FoxO1 I. Quantification of protein level of FoxO3 J. Inflammatory gene mRNA levels were measured by RT-qPCR in BMDM from p21^+/-^ SD, p21^+/-^ HFD, p21^+/Tert^ HFD and p21^+/TertCi^ HFD mice *p<0.05 vs. p21^+/-^ SD mice (white bars), ^#^p<0.05 vs. p21^+/-^ HFD (grey bars). Student’s t test or one-way ANOVA with Fisher multiple comparison test.

### Inactivation of TERT’s catalytic function is associated with only weak metabolic improvements

Our snRNA-seq analysis indicates that TERT and TERT^Ci^ expression exert similar effects on LAMs. To assess whether TERT’s catalytic activity is required for the distinct phenotypes that we previously reported in p21^+/Tert^ obese mice [21], we evaluated these phenotypes in the p21^+/TertCi^ mouse model. In contrast to catalytically active TERT, which we previously reported to improve both glucose intolerance and HFD-induced insulin resistance [21], expression of TERT^Ci^ had only a modest effect on glucose intolerance and failed to improve insulin resistance in HFD-fed mice (Figure 7A-B). This indicates that telomerase catalytic activity plays an important role for mitigating HFD-induced metabolic dysfunction (Figure 7A-B). This is further supported by the lack of differences between p21^+/-^ and p21^+/TertCi^ obese mice in adipocyte size, density within the AT, cell proliferation, and adipogenic gene expression (Figure 7C-G). In line with this, preadipocytes isolated from obese p21^+/TertCi^ mice exhibited mitochondrial dysfunction evidenced by decreased oxygen consumption (Figure 7H-I). suggesting that maintenance of mitochondrial function in ASPCs relies on active telomerase [32]. Similarly, *in vitro* differentiation of ASPCs in mature adipocytes was impaired in both p21^+/-^ and p21^+/TertCi^ mice under HFD compared to pre-adipocytes isolated from SD-fed p21^+/-^ mice (Figure 7J-K). Finally, preadipocytes isolated from from p21^+/TertCi^ obese mice exhibited senescence levels intermediate between those observed in p21^+/-^and p21^+/Tert^ mice (Figure 7L-M).

**Figure 7.**
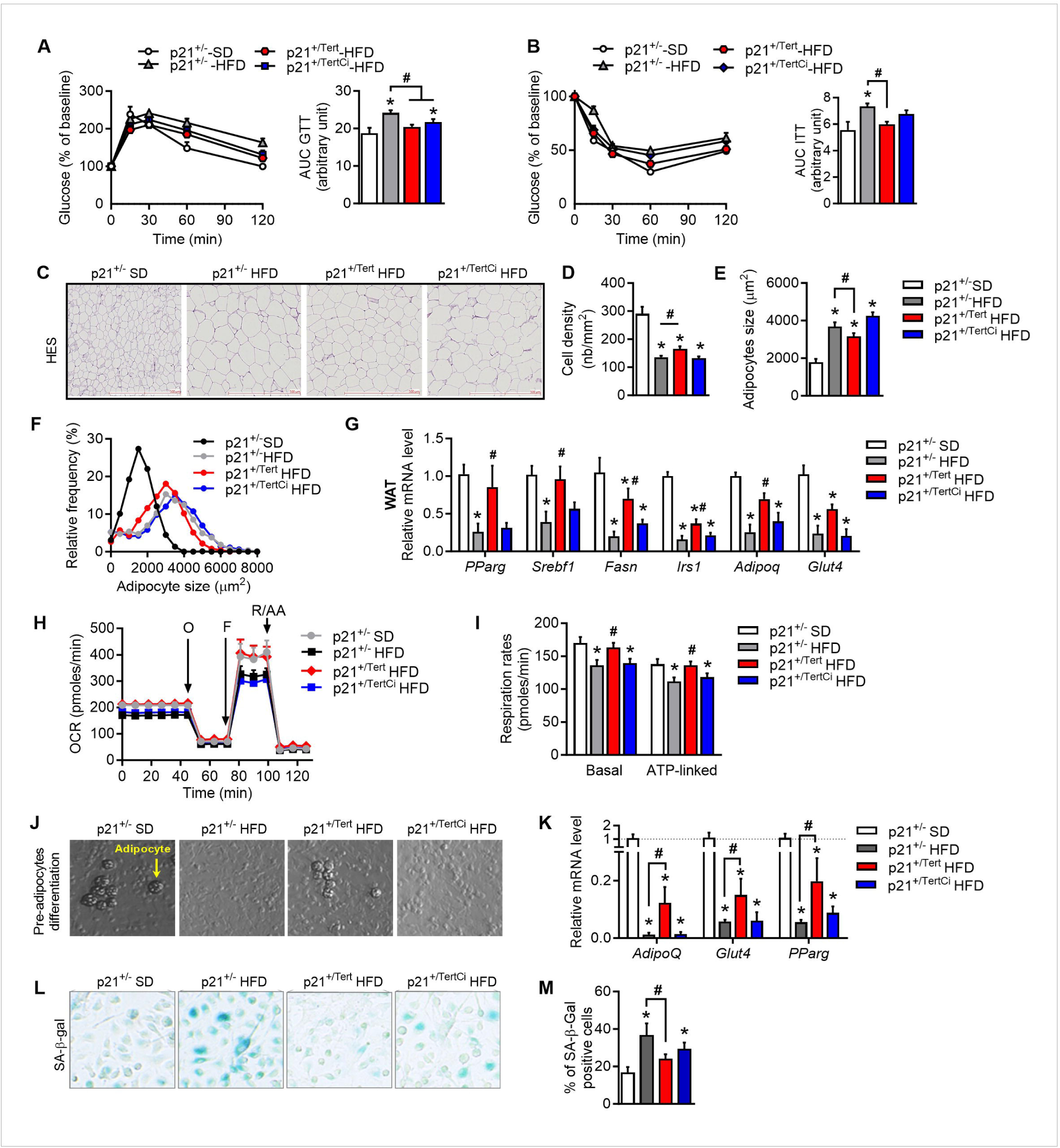
Catalytic inactivation of Tert abolishes its metabolic benefits. A. Left, Glucose Tolerance Test (GTT) was performed by an intraperitoneal injection of glucose (1.5g/kg) and measurement of glycemia via tail clip (Caresens® N, DinnoSanteTM) at different time points. Right, area under the curve (AUC) of the GTT B. Left, Insulin Tolerance Test (ITT) was performed by an intraperitoneal injection of insulin (0.3 UI/kg) and glucose was measured via tail clip (Caresens® N, DinnoSanteTM) at different time points. Right, area under the curve (AUC) of the ITT C. Hematoxylin and eosin (H&E) staining of visceral adipose tissue (WAT) D. Relative frequency of adipocyte sizes in the indicated groups of mice was plotted using GraphPad E. Adipocytes size in the images of H&E-stained WAT was quantified using AdipoSoft ImageJ software F. Adipocyte cell density in the images of H&E-stained WAT was quantified using AdipoSoft ImageJ software G. Expression of the genes involved in lipogenesis, lipolysis and fatty acid oxidation was measured by RT-qPCR in eWAT H. Oxygen consumption rate analyzed by Seahorse in ASPC cell cultures established from WAT I. Respiratory rates calculated based on oxygen consumption data J. Representative photographs showing adipocytes formation from ASPCs in cell culture. Scale is indicated on the image. K. Expression of the genes involved in lipogenesis was measured by RT-qPCR in ASPCs cell cultures established from WAT L. SA-β-gal staining of cultured ASPCs from the WAT. Scale is indicated on the image M. Quantification of SA-β-gal positive cells in the cultures established from WAT of the indicated groups of mice *p<0.05 vs. p21^+/-^ SD mice (white bars), ^#^p<0.05 vs. p21^+/-^ HFD (grey bars). Student’s t test or one-way ANOVA with Fisher multiple comparison test.

## Discussion

We recently reported that p21 promoter-driven TERT expression in HFD-fed obese mice is linked to reduced p21 levels in adipose stem and progenitor cells (ASPCs), preservation of telomere length, lower oxidative DNA damage, and attenuated cellular senescence [21]. These changes collectively promoted the expansion and adipogenic differentiation of ASPCs, leading to the formation of mature adipocytes. We proposed that this coordinated set of effects, particularly the enhancement of adipocyte formation, contributes significantly to the improved insulin sensitivity and reduced glucose intolerance observed in p21^+/Tert^ mice subjected to an HFD [21].

Numerous studies revealed that expression of telomerase has additional roles unrelated to its ability to elongate telomeres [15], in particular in stem cells [33], in progenitor cells [34,35], and also in differentiated cells [36]. We thus wondered whether the effects associated with TERT conditional expression were independent of its catalytic activity. To this end, we expressed a catalytically inactive TERT under the control of the p21 promoter. We found that expression of either TERT or TERT^Ci^ markedly blunted the HFD-induced upregulation of p21, particularly in macrophages, supporting a non-canonical role for TERT in modulating p21 expression. This reduction was accompanied by an attenuation of inflammation-associated gene overexpression within the stromal vascular fraction (SVF). Because p21 expression was particularly elevated in macrophages of HFD-fed control mice, we focused on macrophage dynamics across the three groups of obese mice. In control mice, we observed a strong expansion of MPCs driven largely by the emergence of LAM clusters, especially Trem2+ and Gdf15+ macrophages. In contrast, both p21^+/Tert^ and p21^+/TertCi^ mice displayed a markedly reduced expansion of these macrophage clusters. Although Gdf15+ macrophages increased modestly in these models, the Trem2+ population was nearly absent consistent with a markedly reduced Trem2 transcriptomic signature in both p21^+/Tert^ and p21^+/TertCi^ macrophages. This reduction in the two LAM clusters was accompanied by an increased abundance of resident-tissue macrophages (RTM) in both p21^+/Tert^ and p21^+/TertCi^ obese mice. These findings suggest that TERT expression may either impair the differentiation of monocytes and resident macrophages into LAMs or reduce monocyte recruitment into AT.

These results raise the question of whether the differences in macrophage population distribution in the AT of obese mice come from an intrinsic alteration of macrophages linked to TERT (or TERT^Ci^) expression, or from a disruption in cell-cell communication, particularly between ASPCs/adipocytes and macrophages [37]. We found that ASPCs/adipocytes engaged extensively with LAMs, as reflected by numerous ligand-receptor pairs. Strikingly, p21^+/Tert^ and p21^+/TertCi^ mice had reduced ASPCs/LAM communication involving C3, Csf1, Fgf1, Fgf7, Tgfb2, and Semad3, molecules typically linked to inflammation and tissue remodelling. Notably, it was previously reported that C3aR expression increases upon HFD in adipocytes and macrophages of the AT of obese mice, and that loss of C3aR protects mice from early diet-induced obesity, improves insulin sensitivity, reduces liver steatosis, and diminishes adipose inflammation by lowering macrophage infiltration and pro-inflammatory cytokine production [38]. Along the same line, CSF1 is broadly expressed across mouse tissues predominantly by mesenchymal lineage cells and its local production is essential for sustaining tissue-specific macrophage populations [7,39]. Although the CSF1/CSF1R axis is known to support macrophage survival and maintenance, a direct role in the LAM lineage has yet to be demonstrated. Our findings suggest that CSF1 produced by adipocytes and/or ASPCs may create a niche that promotes recruited monocytes or resident macrophages to differentiate and persist as specific LAMs, a process that appears to be counteracted in p21^+/Tert^ and p21^+/TertCi^ AT. On its side, FGF2 was shown to increase in the AT of HFD-fed obese mice, and recombinant FGF2 enhanced NLRP3 inflammasome activation and pro-inflammatory signalling in adipocytes, supporting its role as an adipokine that exacerbates adipose inflammation [40]. These reduced interactions in the TERT-expressing models aligns with their reduced inflammatory profile.

To further support our model of ASPCs/macrophage communication, we isolated ASPCs from p21^+/Tert^ mice and analysed the proteins released in their secretome. We found that p21^+/Tert^ ASPCs secreted higher levels of proteins involved in vesicle trafficking, endosomal maturation, endolysosomal regulation and cargo sorting. The enrichment of proteins such as Rab21, Rab5a, Sec22b, Sec23b, Vps16, and SNX family members, none of which possess signal peptides and are not secreted through the classical ER–Golgi pathway, strongly indicates that p21^+/Tert^ ASPCs release vesicular trafficking components via Golgi-bypass, non-conventional secretion mechanisms (e.g., EV-mediated or autophagy-linked secretion [41]. Conversely, the pronounced depletion of canonical secreted proteins (ECM proteins, MMPs, TIMPs, integrins, cadherins, cytokines, and growth factors, classical secretory cargoes requiring signal peptides [42] suggests a broad reduction or dysfunction of the conventional ER–Golgi secretory pathway in these cells. Together, these findings indicate a shift toward unconventional protein secretion in p21^+/Tert^ ASPCs. Of note, it was shown recently that senescent cells were particularly sensitive to the inhibition of their secretory apparatus [43]. Consistent with our snRNA analyses, we observed that C3 and Csf1, and also Ccl2, which regulate either macrophage activity, or monocyte/macrophage survival, differentiation, and recruitment (see above) were underrepresented in the supernatant of untreated p21^+/Tert^ ASPCs. These proteins are normally secreted though the conventional secretory pathway, although we cannot rule out the possibility that their reduced abundance reflect lower expression in p21^+/Tert^ ASPCs. In line with the results obtained with untreated cells, IFN-γ/TNF-α treated p21^+/Tert^ ASPCs displayed reduced levels of multiple inflammatory and chemoattractant cytokines (Ccl2, Ccl5, Ccl7, Cxcl1, Cxcl5, Cxcl9, Cxcl10 and Vegfa), all of which are secreted though the conventional ER/Golgi pathway, and includes several well-established SASP components.

These results reinforce the notion that the altered macrophage landscape in the AT of p21^+/Tert^ (and p21^+/TertCi^) mice arises, at least in part, from changes in ASPC-macrophage interactions driven by ASPC reprogramming and corresponding alterations in their secretory machinery. Of note, a recent study showed that senescent cells are particularly sensitive to inhibition of the conventional ER/Golgi/vesicular trafficking pathway [43]. Whether the shift in expression and/or secretion we observe reflects the reduced senescence we recently reported in p21^+/Tert^ ASPCs [21] remains to be determined.

Interestingly, we found that Pparγ expression was markedly reduced in macrophages from both p21^+/Tert^ and p21^+/TertCi^ mice. Pparγ is a central transcriptional regulator driving the differentiation of macrophages toward LAM. Notably, LAM with high PPARγ activity have been shown to support adipose tissue expansion during high-fat diet feeding [44]. Because PPARγ functions as a lipid-sensing nuclear receptor activated by fatty acids and oxidized lipid species, its low expression in these mice suggests a diminished capacity of macrophages to engage lipid-driven LAM differentiation [45]. Our results raise the possibility that adipocytes in p21^+/Tert^ and p21^+/TertCi^ mice have a better lipid-buffering efficiency, thereby reducing lipid spillover into the stromal-vascular niche. This attenuation of extracellular lipid availability would, in turn, limit the activation of PPARγ-dependent LAM programs and ultimately restrain LAM expansion. Importantly, our single-cell data also revealed a concomitant reduction in Nr1h3 (encoding LXRa), another master regulator of lipid handling transcriptional programs in LAM [4,46]. Therefore, the dual reduction of Pparγ and Nr1h3 points to a broader impairment in the macrophage lipid-sensing axis, rather than a defect restricted to a single pathway. Because LXRa and PPARγ act cooperatively to drive lipid-dependent reprogramming and stabilize the LAM phenotype, reduced Nr1h3 expression likely further compromises the ability of macrophages to transition into a lipid-adapted, LAM-like state.

Interestingly, macrophages, as do ASPCs, from obese p21^+/Tert^ (and p21^+/TertCi^) mice show increased AKT activation. This is reflected transcriptionally by the down-regulation of inflammatory genes such as IL6 and TNFα in BMDMs from obese p21^+/Tert^ and p21^+/TertCi^ mice. This pattern is consistent with reports indicating that reduced AKT activity drives pro-inflammatory macrophage polarization [30,31]. Together, these findings indicate that TERT independently of its catalytic activity may enhance AKT signalling in specific contexts such as obesity.

One important unresolved question is whether AT LAMs are adaptive or maladaptive. Previous studies support both possibilities [47]. Trem2 silencing in myeloid cells prevents LAM accumulation in adipose and improves metabolic function [4]. While transgenic Trem2 overexpression aggravates diet-induced obesity and metabolic dysfunction [48], and adoptive transfer of CD9-positive LAMs triggers adipose tissue inflammation [8], together suggesting LAMs may have early buffering effects but later become harmful. Consistent with dual adaptive-maladaptive properties, LAMs co-express genes with anti-inflammatory, pro-inflammatory, and pro-fibrotic functions, with certain subpopulations exhibiting higher pro-inflammatory signatures [8,47,49–51]. Taken together, this suggests the reduction in LAMs in TERT could dampen an inflammation, inflammation-resolution, and fibrosis cycle that improves overall homeostasis in obese AT.

In this study, we show that both TERT and TERT^Ci^ reduce the number of p21^high^ macrophages and Gdf15- and Trem2-expressing macrophages. This finding raises the question of whether TERT^Ci^ expression provides metabolic benefits comparable of those of TERT in the context of obesity-associated disorders in mice. Our *in vivo* and *in vitro* data indicate that the formation of functional adipocytes is primarily enhanced by TERT, but not by TERT^Ci^, suggesting that TERT’s catalytic activity is essential for the generation of mature adipocytes. Accordingly, the maintenance of functional telomeres appears to be required for the steps leading to adipocyte maturation. These observations are consistent with the greater ability of TERT, relative to TERT^Ci^, to improve insulin-resistance in obese mice. TERT’s non-canonical functions, including its capacity to reduce Gdf15 and Trem2 expressing LAMs, may contribute to its metabolic effects. However, these non-canonical functions alone may be necessary but not sufficient to fully ameliorate the metabolic defects induced by a high-fat diet.

## Materials and Methods

### Animals

Mice p21^+/-^, p21^+/Tert^ and p21^+/TertCi^ were housed under controlled conditions of temperature (21±1°C), hygrometry (60±10%) and lighting (12 h per day). Animals were acclimatized in the laboratory for one week before the start of the experiments. Mice were fed either a standard diet SD (A04, SAFE Diet, Augy, France) or a high fat diet HFD (60 kcal % fat, SAFE Diet, Augy, France). All animals received care according to institutional guidelines, and all experiments were approved by the Institutional Ethics committee number 16, Paris, France (licence number 16-090). During follow-up, animals underwent body-weight, metabolic assessments. Mice were euthanatized and organs and blood were collected and processed for further evaluations.

### Analysis of mRNA expression

For RNA extraction the samples were lysed with Qiazol (Qiagen, France) in the presence of chloroform, and total RNA was purified on mini-columns using Rneasy extraction kit (Qiagen, ref 74104). First-strand cDNA was synthesized from total RNA using the High-Capacity cDNA Reverse Transcription Kit (Life Technologies, ref 4368814). Quantitative real-time PCR (qPCR) was performed in a StepOnePlus Real-Time PCR. Gene expression was assessed by the comparative CT (ΔΔCT) method with β-actin as the reference gene.

### Histology and immunohistochemistry

Fresh visceral adipose tissue was fixed in 10% phosphate-buffered formalin overnight. Paraffin wax sections of 5 µm were prepared for immunostaining (haematoxylin-eosin staining).

### Imaging flow cytometry analysis of the SVF

Purified SVF cells were incubated with a panel of antibodies (CD45 BD Bioscience 561047, CD31 BD Bioscience 612802, CD34 BioLegend 119314, CD29 BioLegend-102218, ScaI BioLegend 108127, CD24 BioLegend-101823, PDGFRα BioLegend-135905, F4/80 BioLegend-123115, CD11b BD Bioscience-565976, CD11c BD Bioscience-749038, Live Dead Nir Cytek R7-60008) for 30 min on ice. Cells were then washed with 0.5% FBS/PBS FACS buffer and resuspended in 300 µL of FACS buffer. Cells were detected and their fluorescence was measured using Aurora flow cytometer (Cytek, Amsterdam, Netherland). Data analysis was performed using SpectroFlow and OMIQ softwares.

### Cellular Bioenergetic Analysis Using the Seahorse Bioscience XF Analyzer

Bioenergetic profiles of the adipocytes were determined using a Seahorse Bioscience XF24 Analyzer (Billerica, MA, USA) that provides real-time measurements of oxygen consumption rate (OCR), indicative of mitochondrial respiration, and extracellular acidification rate (ECAR), an index of glycolysis as previously described [52].

### RNA-Seq and Data Analysis

Quality control of samples. RNA sequencing was performed by Novogene company (Cambridge, UK). RNA integrity was assessed using the RNA Nano 6000 Assay Kit of the Bioanalyzer 2100 system (Agilent Technologies, CA, USA). Library preparation for Transcriptome sequencing. Total RNA was used as input material for the library preparations. Briefly, mRNA was purified from total RNA using poly-T oligo-attached magnetic beads. Fragmentation was carried out using divalent cations under elevated temperature in First Strand Synthesis Reaction Buffer (5X). First strand cDNA was synthesized using random hexamer primer and M-MuLV Reverse Transcriptase (Rnase H-). Second strand cDNA synthesis was subsequently performed using DNA Polymerase I and Rnase H. Remaining overhangs were converted into blunt ends via exonuclease/polymerase treatment. After adenylation of the 3’ ends of DNA fragments, Adaptor with hairpin loop structure was ligated to prepare for hybridization. In order to select cDNA fragments in the reange of 370-420 bp, the library fragments were purified with AMPure XP system (Beckman Coulter, Beverly, USA). Then PCR was performed with Phusion High-Fidelity DNA polymerase, using the Universal and Index (X) primers. At last, PCR products were purified (AMPure XP system) and library quality was assessed on the Agilent Bioanalyzer 2100 system. Clustering and sequencing. The clustering of the index-coded samples was performed on a cBot Cluster Generation System using TruSeq PE Cluster Kit v3-cBot-HS (Illumia) according to the manufacturer’s instructions. After cluster generation, the library preparations were sequenced on the Illumina Novaseq platform generating 150 bp paired-end reads. Sequencing quality control was determined using the FastQC tool (http://www.bioinformatics.babraham.ac.uk/projects/fastqc/) and aggregated across samples using MultiQC (v1.7) [29]. Reads with a Phred quality score of less than 30 were filtered out. Reads were mapped to a customized mouse mm10 genome (containing the mCherry transgene) using Subread-align (v1.6.4) [54] with default parameters. Gene expression was determined by counting mapped tags at gene levels using featureCounts (v1.6.4) [55], and differentially expressed genes (DEG) were identified using DESeq2 (v1.26.0) [56]. Statistical significance was inferred at P<0.05 (Benjamini-Hochberg corrected). Gene Ontology (GO) enrichment analysis was performed on DEGs using clusterProfiler (v4.6.0) [57] and considering a background. Only the biological process (BP) category was chosen and GO terms with corrected P value less than 0.05 were considered significantly enriched. Enrichment heatmaps were generated using heatmap.2 (v<u>3.1.3.1)</u> [58], highlighting the 10 most differentially regulated genes per GO term. String analysis (https://string-db.org) was performed to investigate interactions between differentially expressed gene.

### Single-nuclei RNAs analyses

The snRNAseq analysis was conducted following the same methodology as previously described [21] with the additional step of Doublet removal using DoubletFinder [59]. The scRNAseq analysis of good quality cells was performed in R v4.4.0 using Seurat Package v5.0.3 [60]. After normalization, we identified the top 2000 variable genes for each library and the libraries were integrated using the RPCA method. For all the integrated objects, we performed linear dimensional reduction (PCA), cell clustering and data visualization using UMAP and t-SNE. Differentially expressed genes in the MPC cluster were calculated using a Wilcoxon Rank Sum test in Seurat. Additionally, LAM signature were calculated using the same methodology used in Rizzo et al. [26]. Intercellular communication was inferred using the CellChat R package (v1.6.2) [37] and its ligand-receptor database. Communication probabilities between cell types were computed using the mass action model based on the expression of over-expressed ligand-receptor pairs. The analysis quantified overall information flow, determined functional similarity via hierarchical clustering, and identified key signalling roles (e.g., sender, receiver) for each cell type. Single-cell trajectory and pseudotime analysis was performed using the Monocle 2 R package (v2.32.0) [28] to reconstruct cellular differentiation paths. A Cell Data Set was created using the MPC subclustering. The trajectory was learned using the Discriminative Dimensionality Reduction (DDRTree) algorithm, allowing cells to be ordered by pseudotime from a not defined root.

### Bone Marrow Derived Macrophages (BMDM) cell culture

Bone marrow cells were isolated from the tibia and femur of p21+/- SD, p21+/- HFD, p21+/Tert HFD, and p21+/TertCi HFD mice. The cells were cultured in high-glucose DMEM supplemented with 10% fetal bovine serum (FBS) (Merck), 100 U/ml penicillin, 10 mg/ml streptomycin, and 25 ng/ml recombinant mouse M-CSF (macrophage colony-stimulating factor) to induce differentiation. Following differentiation, cells were harvested for subsequent analyses.

### Telomere Shortest Length Assay (TeSLA)

TeSLA (Telomere Shortest Length Assay) was performed as previously described [21] to measure the distribution of the shortest telomeres in cells.

### Western Blot Analysis

BMDM cells were lysed in cell lysis buffer (Cell Signaling, Danvers, MA France) supplemented with 1% phenylmethylsulfonyl fluoride (PMSF). Protein samples were resolved on 12% bis-Tris gels followed by transfer to nitrocellulose membrane. Antibodies for total Akt (#4991), phospho-Akt (Ser473) #4060), Foxo1 (#2880), Foxo3a (#9467) were from Cell Signaling and antibodies for β-actin (sc-47778) were from Santa Cruz. Bands were visualized by enhanced chemiluminescence and quantified using ImageJ software.

### Isolation and culture of ASPCs

Visceral fat was excised from the mice, minced with scissors, and digested for one hour at 37°C in 1 mg/ml collagenase type II digestion buffer (Life Technology, France; 17100017) in sterile Hank’s Balanced Salt Solution (Life Technologies, ref 14185-052) containing 3% bovine serum albumin. After digestion, stromal vascular fraction (SVF) containing ASPCs (adipose stem/progenitor cells) were separated from adipocytes by centrifugation (300g, 3min) and two filtration steps (70µm and 40µm cell strainers). SVF were plated in a 6-well plate in DMEM supplemented with 0.5% Pen/Strep, 10% new born calf serum and 1% pyruvate.

### Treatment of ASPCs with IFN-γ and TNF-α

Adipose stem and progenitor cells (ASPCs) were isolated from visceral adipose tissue and cultured until reaching 70–80% confluence. ASPCs were then treated with recombinant mouse TNF-α (10 ng/mL) (Biotechne, ref 410-MT) and IFN-γ (20 ng/mL) (Biotechne, ref 485-MI) for 36 hours. Following stimulation, cells were serum-starved for 24 hours collecting conditioned supernatants and subsequent proteomic analysis.

### Proteomics

The resulting cleared supernatants were further dialyzed and concentrated against 100 mM ammonium bicarbonate using Amicon4 Ultra, 3Kd MWCO (Millipore). Secretome extracts were dried using a SpeedVac vacuum concentrator, and each protein extract was stacked on NuPAGE™ in a single band to perform trypsin digestion before mass spectrometry analysis using an Orbitrap Fusion Lumos Tribrid (ThermoFisher Scientific, San Jose, CA) in data-independent acquisition (DIA) mode. Protein identification and quantification were processed using the DIA-NN 1.8 algorithm [61] and DIAgui package (https://github.com/marseille-proteomique/DIAgui) [62]. The statistical analysis was done with the Perseus program (version 1.6.15.0) [63]. Differential proteins were detected using a two-sample t-test at 0.05 permutation-based false discovery rate. Statistical analysis was performed using the standard two-tailed Student’s t-test, and P-value < 0.05 was considered significant.

### Statistical analysis

Data are expressed as mean values ± standard error of the mean (SEM). Statistical significance was tested using either one or two-way analysis of variance (ANOVA) with Fisher multiple comparison test. The results were considered significant if the p-value was <0.05.

### Data availability

Single-nucleus RNA-seq data has been deposited in the Gene Expression Omnibus (GSE261439).

## Acknowledgements

We thank Jean-Charles Graziano from CRCM’s animal facility, Manon Richaud from CRCM Cytometry platform and the CRCM’s Integrative BioInformatics platform (Cibi). We would like to thank Frédéric Fiore (CIPHE, Marseille) for his advice on generating mouse models. We thank Stoyan Ivanov (LP2M, Nice) and Thierry Galli (IPNP, Paris) for helpful discussions. Work in VG’s Lab was supported by “La Ligue Nationale Contre le Cancer”, Equipe Labellisée, the “Institut National du Cancer” (INCA, PLBIO 2019), the Agence Nationale de la Recherche (ANR) (Grant THALATEL), the cross-cutting Inserm program on aging (AGEMED) and the Inserm INTERAGING Program. This study was partly supported by research funding from Gefluc and Canceropole Sud. Core support from MRC (MC_U120085810) and CRUK (C15075/A28647) funded this research in J.G’s laboratory. W.S. was funded by the MRC (MC_UP1605/7) and Wellcome Trust (219602/Z/19/Z). MV and JMB were financed by INSERM.

## Author Contributions

L.B. and V.G. designed the study, supervised the project and analyzed the data. L.B coordinated most the experiments and data analysis. L.B, M.B., M.R performed metabolic tests, qPCR, FACS experiments. M.B. and D.C. performed TeSLA experiments. W.S, A.M. and J.G. performed the snRNseq experiment. A.N. and D.L. analyzed snRNAseq data. J.V. analyzed the RNAseq data. J-M.B., S.A. and E.B. performed the proteomic experiment. V.G., L.B., C.L and J.G secured funding. L.B and V.G. wrote the manuscript with inputs of W.S., J.G., D.L., A.N. L.B and V.G are the guarantors of this work and, as such, have full access to all the data in this study and take responsibility for the integrity of the data and the accuracy of the data analysis.

## Supplemental Figure Legends

**Suppl Fig 1.**
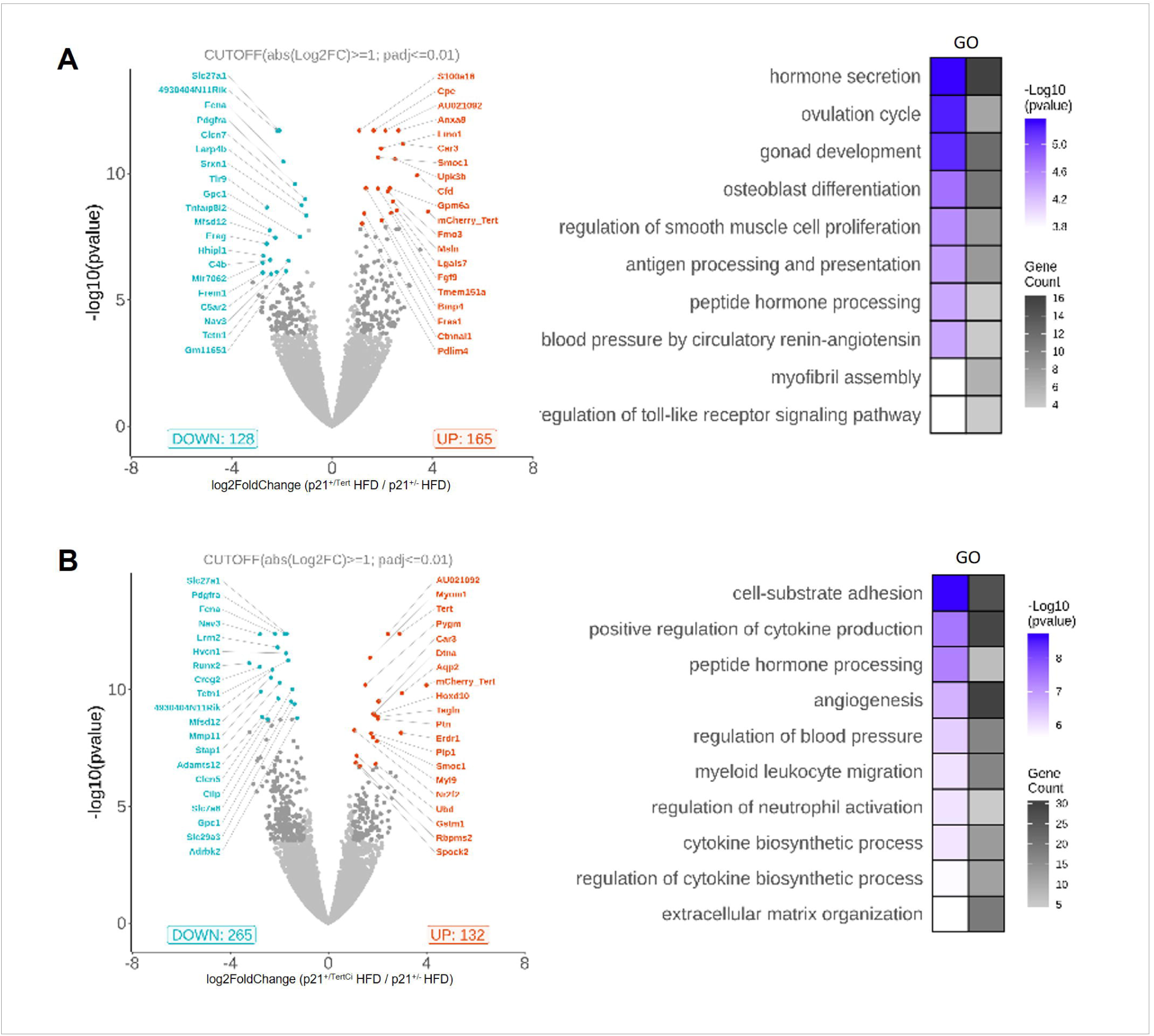
Volcano plots of differentially expressed genes and associated GO analyses. A. Volcano plots indicating the number of significantly (<0.05) differentially expressed between p21^+/-^HFD and p21^+/Tert^ HFD mice and associated GO analysis B. Volcano plots indicating the number of significantly (<0.05) differentially expressed between p21^+/-^HFD and p21^+/TertCi^ HFD mice and associated GO analysis

**Suppl Fig 2.**
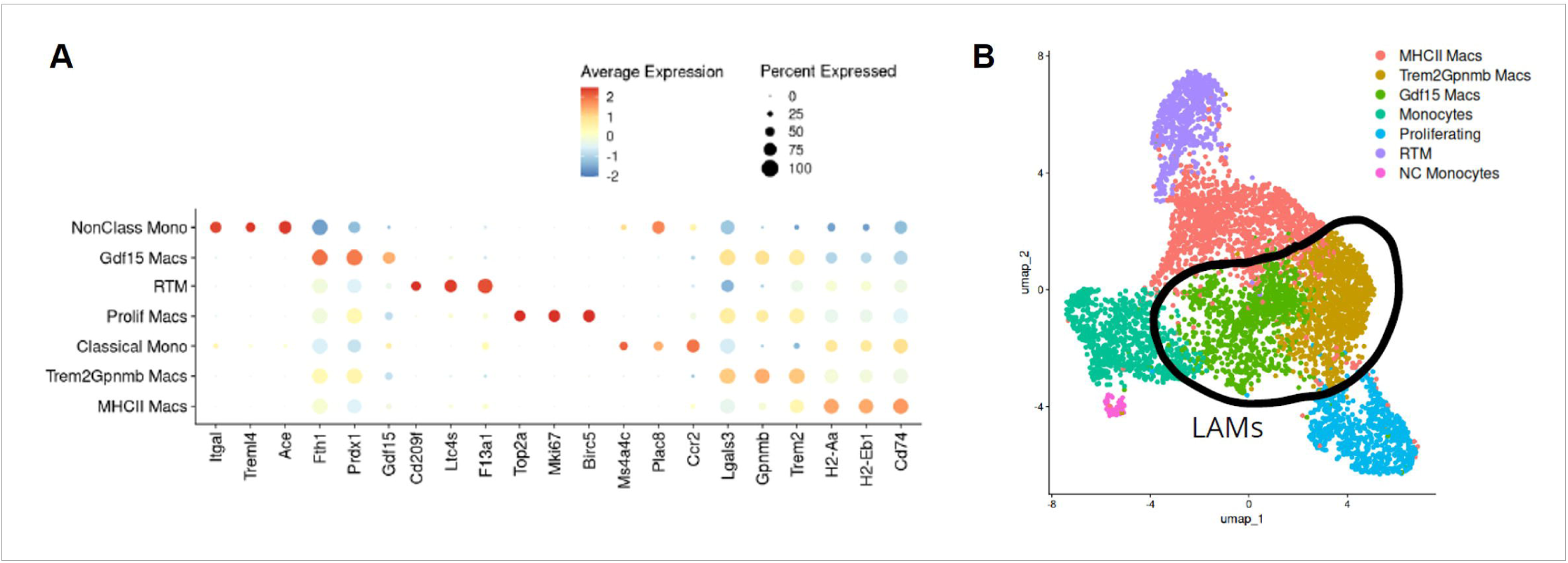
High-resolution annotation of macrophage subclusters. A. Top unbiased markers of the different ASPC subtypes. Dotplots represent scaled expression per gene. B. UMAP identifying the different ASPC subclusters based on the unbiased markers. Differentially expressed genes in the MPC cluster were calculated using a Wilcoxon Rank Sum test in Seurat.

**Suppl Fig 3.**
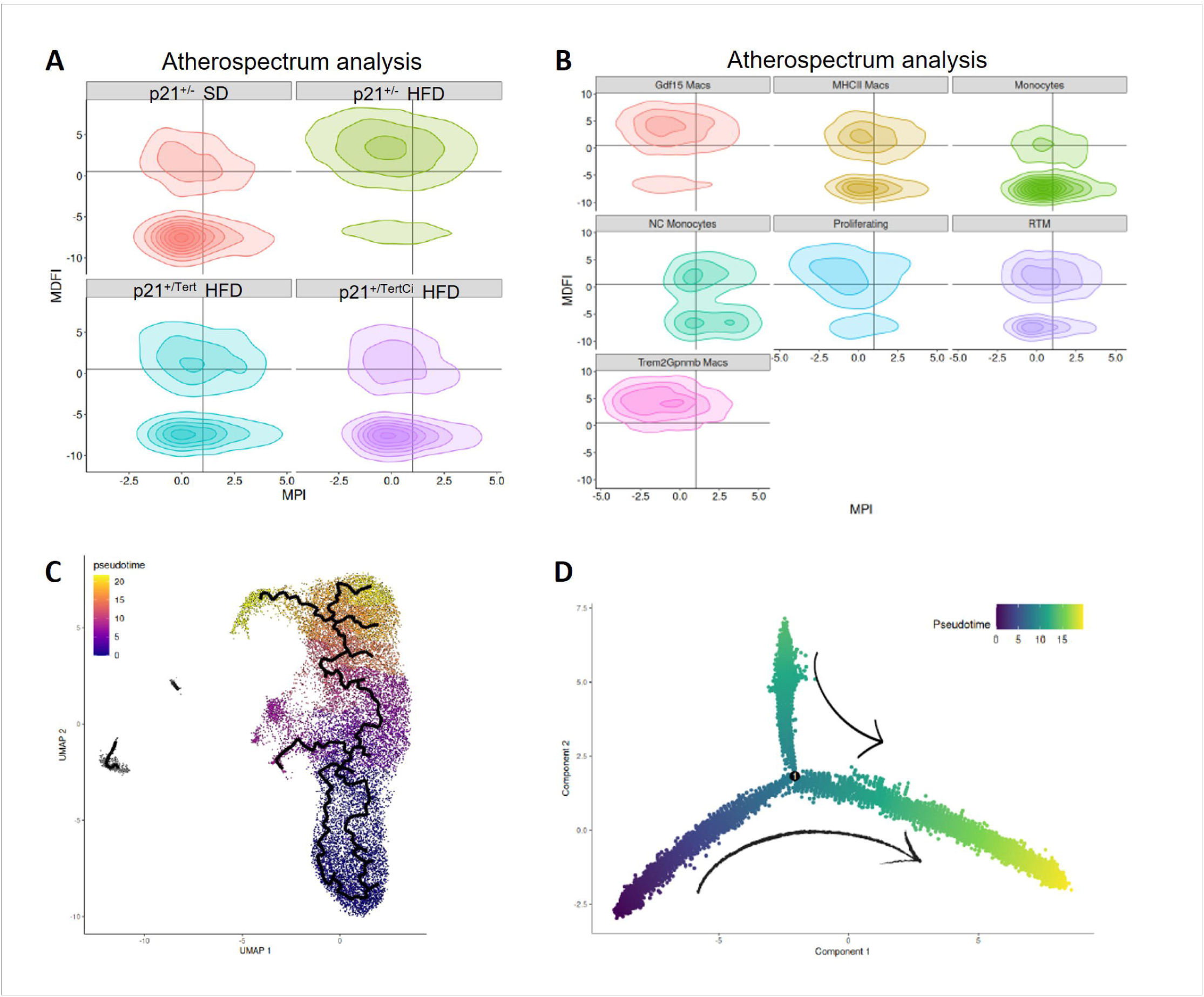
Atherosclerum and single-cell trajectory analysis. A. Atherosclerum analysis between the different group of mice B. Atherosclerum analysis between the different ASPC subclusters C. Single-cell trajectory and pseudotime analysis was performed using the Monocle 2 R package (v2.32.0) [28] to reconstruct cellular differentiation paths. D. Trajectory analysis across the identified macrophage subclusters

**Suppl Fig 4.**
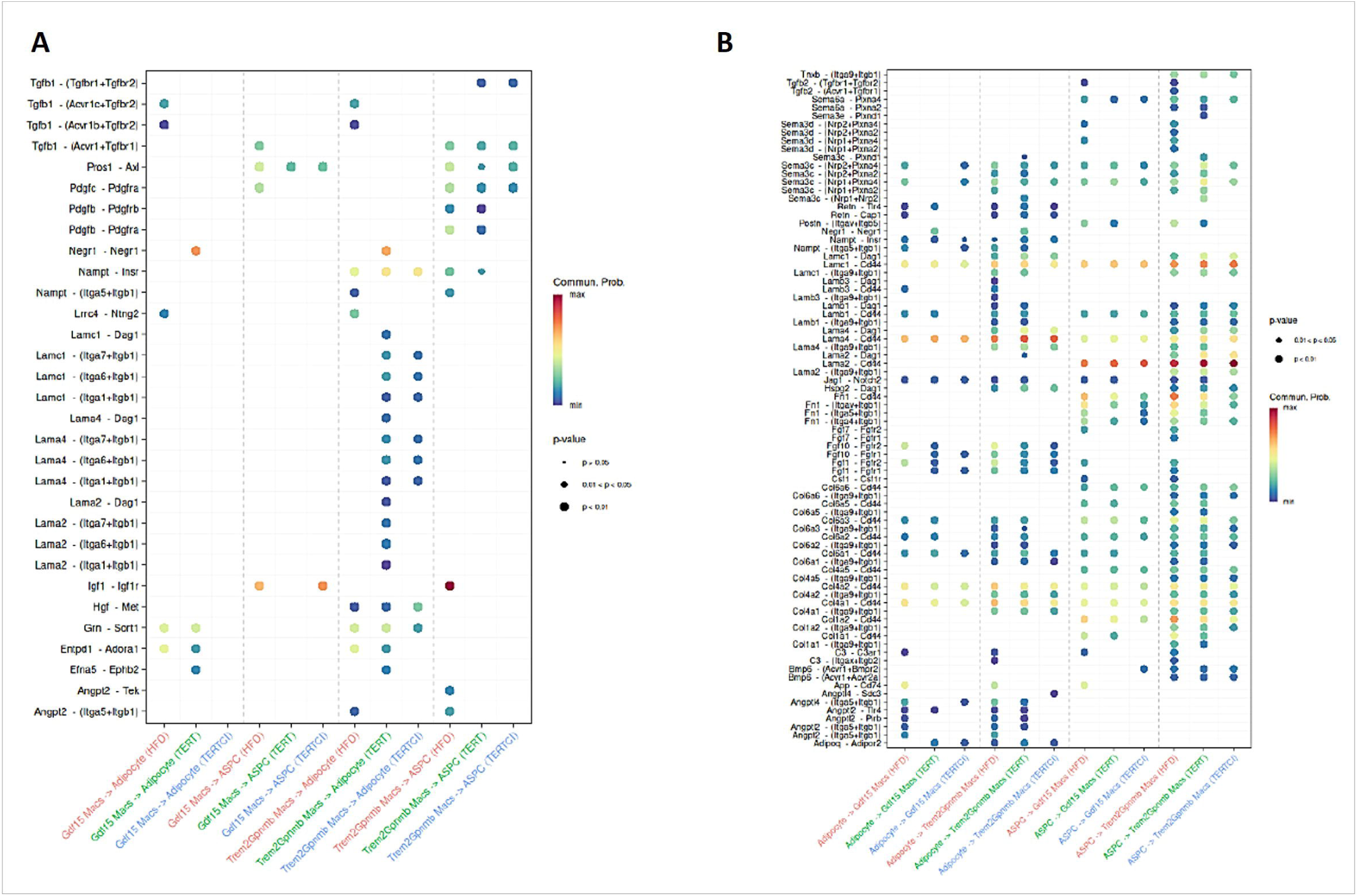
Communication probabilities between ASPC-derived adipocytes and macrophages in adipose tissue (AT) from obese p21^+/-^, p21^+/Tert^ and p21^+/TertCi^ obese mice. A. Communication probabilities from Trem2+ and Gdf15+ macrophages to ASPCs and adipocytes. B. Communication probabilities from ASPCs and adipocytes to Trem2+ and Gdf15+ macrophages. Communication probabilities between cell types were computed using the mass action model based on the expression of over-expressed ligand-receptor pairs.

**Suppl Fig 5.**
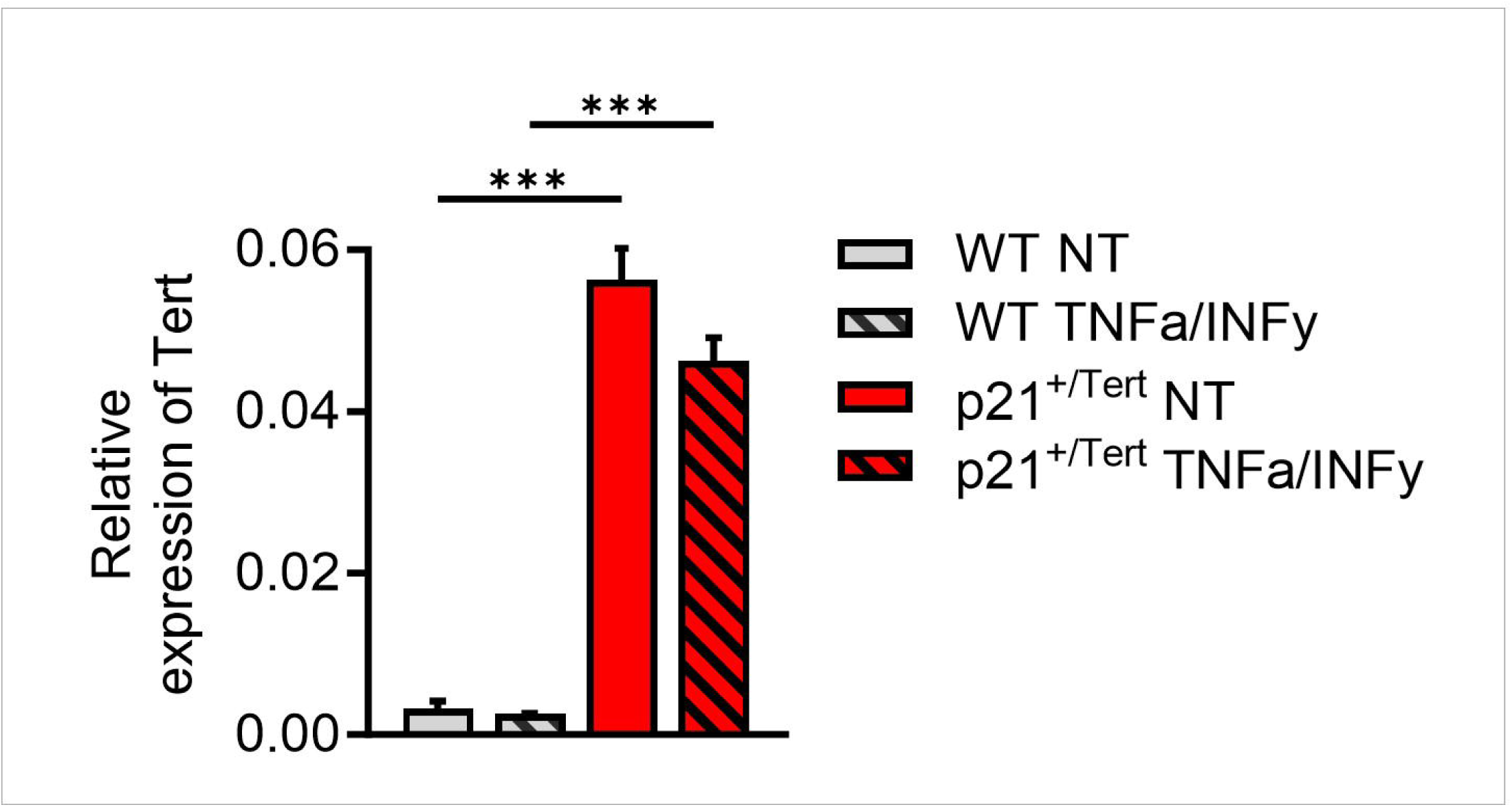
TERT expression in WT and p21^+/Tert^ ASPCs, with or without IFN-γ and TNF-α treatment. WT and p21^+/Tert^ ASPCs were treated or not (NT) with IFN-γ and TNF-α *in vitro*. Expression levels of Tert were measured by RT-PCR from cell pellets of the indicated ASPCs. *p<0.05. one-way ANOVA with Fisher multiple comparison test.

## Dataset Legends

**Dataset 1.** RNAseq of SVF of p21^+/-^ SD, p21^+/-^ HFD, p21^+/Tert^ HFD, p21^+/TertCi^ HFD

**Dataset 2.** Number of macrophages for each subcluter and for each condition

**Dataset 3.** Differentially Expressed Genes (DEGs) for the MPC cluster across the individual conditions

**Dataset 4.** Proteomic analysis of secreted protein between WT and p21^+/Tert^ untreated APSCs

